# Characterising phagocytes and measuring phagocytosis from live *Galleria mellonella* larvae

**DOI:** 10.1101/2023.10.09.558207

**Authors:** Jennie S. Campbell, Attila Bebes, Arnab Pradhan, Raif Yuecel, Al Brown, James G. Wakefield

**Affiliations:** Living Systems Institute, University of Exeter, Exeter, UK; Exeter Centre for Cytomics, Henry Wellcome Building for Biocatalysis, Biosciences, University of Exeter, Exeter, UK; Medical Research Council Centre for Medical Mycology, University of Exeter, School of Biosciences, Geoffrey Pope Building, Exeter, UK

## Abstract

Over the last 20 years, the larva of the greater waxmoth, *Galleria mellonella*, has rapidly increased in popularity as an *in vivo* mammalian replacement model organism for the study of human pathogens. Despite this, experimental readouts of response to infection are generally limited to observing the melanisation cascade – where the organism turns black as part of the systemic immune response – and quantifying larval death over time. As an invertebrate, *Galleria* harbour an innate immune system comprised of both humoral components and a repertoire of innate immune cells – termed hemocytes. Though information on subtypes of hemocytes exist, there are conflicting reports on their exact number and function. Flow cytometry has previously been used to assay *Galleria* hemocytes, but protocols include both centrifugation and fixation - physical methods which have the potential to affect hemocyte morphology prior to analysis. Here, we present a method for live hemocyte analysis by flow cytometry, revealing that *Galleria* hemocytes constitute only a single resolvable population, based on relative size or internal complexity. Using fluorescent zymosan particles, we extend our method to show that up to 80% of the *Galleria* hemocyte population display phagocytic capability. Finally, we demonstrate that the developed assay reliably replicates *in vitro* data, showing that cell wall β-1,3-glucan masking by *Candida albicans* subverts phagocytic responses. As such, our method provides a new tool with which to rapidly assess phagocytosis and understand live infection dynamics in *Galleria*.

## Introduction

Larvae of the greater waxmoth, *Galleria mellonella*, are rapidly gaining popularity as the mammalian replacement model organism of choice to study a variety human pathogens to which they are susceptible [1–8]. This is due both to their relatively large size – up to 2.5 cm [9] – which makes them amenable to handling and accurate dosing, and to their viability at 37 °C (i.e. human physiological temperature).

As an invertebrate, *Galleria mellonella* is equipped with an innate immune system – which, like that of mammals, consists of both humoral and cellular responses. A major humoral immune response is the induction of the melanisation cascade. This cascade, which is initiated by the presence of both microbial compounds [10] as well as physiological stressors, cause *Galleria* larvae to turn black following the activation of phenoloxidase, and the melanin produced then aids in the trapping and killing of pathogens [11]. This phenotypic change from a healthy cream colour to a darker brown/black can be scored for simple health readouts from infection experiments [12]. However, whilst this provides a health status readout for the organism, the response cannot be correlated to an analogous mammalian infection response.

In addition to melanisation, *Galleria* larvae generate an array of antimicrobial peptides (AMPs) to combat infection [13,14]. Insect AMPs are primarily secreted from both the fat body [15], which is the equivalent to the mammalian liver and adipose tissue [16], and the cells of the insect immune system. Due to a distinct lack of experimental tools - such as antibodies - available for *Galleria* research, the levels of AMPs cannot be quantified and compared via biochemical means such as Western Blotting. Thus, more laborious techniques including rt-/q-PCR and comparative proteomics/transcriptomics are currently used to understand the humoral responses of *Galleria* larvae to infections and physiological stressors [17–19].

The cellular component of the *Galleria* immune system are innate immune cells - termed hemocytes – found within the hemolymph that circulates through the hemocoel. Hemocytes perform three major immune functions – phagocytosis, encapsulation and induction of the melanisation cascade [15,20,21]. *Galleria* hemocytes were first observed and characterised in 1977– with five cell subtypes being identified [22]. Of these, plasmatocytes and granulocytes (the most abundant) display phagocytic capability *in vitro* [21]. The phagocytic rate of *Galleria* hemocytes has generally been investigated by phase-contrast imaging and manual counting [23–25]. Flow cytometry, which has been used extensively in mammalian immune cell biology to help understand distinct immune cell types and functions, has been applied to *Galleria*, but flow plots vary widely across publications and methods use generally fixed, rather than live, immune cells [26–29]. Though fixation is vital for the detection of intercellular antigens via antibody staining, and for the attenuation of hazardous biological materials, fixatives have been shown to affect both relative cell size and granularity when analysed by flow cytometry [30]. Thus, the use of fixative in the absence of meaningful antigen detection by antibody staining may not be entirely appropriate when investigating infection dynamics and phagocytosis.

Here, we present a method for the analysis of live *Galleria* hemocytes by flow cytometry with minimal sample processing. We demonstrate its efficacy in studying the phagocytosis of fluorescent particles and pathogens following injection into *Galleria* larvae and their subsequent incubation – meaning phagocytosis can be quantified over time. Finally, we show that this *in vivo* phagocytosis assay can detect the subversion of phagocytosis following β-glucan masking by *Candida albicans* grown on alternative carbon sources which recapitulates *in vitro* cell culture data [31].

## Materials and methods

### Galleria mellonella rearing

An in-house colony of *Galleria mellonella* was utilised for all experiments. All stages of the lifecycle were maintained in constant darkness at 30 °C in a temperature-controlled incubator (LEEC). Briefly, fertilised embryos were collected from egg papers placed within adult rearing jars, and were put into diet made according to diet recipe 3 from Jorjão et al [32]. Larvae were allowed to develop undisturbed until needed for experiments.

### Larval injection

Last instar larvae were selected from larval feeding jars for use in experiments. Larvae between 250-350 mg were chosen, and those which showed either melanisation or the presence of a bright dorsal ecdysial line [9] were omitted due to unsuitability for experiments. For injection, larvae were then held over a 1000 μL pipette tip, ventral side up and 10 μL of injectate mixture was injected using a Hamilton syringe (700 series – Merck) connected to a PB600 repeating syringe dispenser through the last right proleg for consistency. Following injection, larvae were kept within petridishes according to the timepoint for analysis. Injected larvae were returned to temperature-controlled incubators as the time course began.

For control injections, *Galleria* larvae were injected with insect physiological saline (IPS – 150 mM sodium chloride, 5 mM potassium chloride, 10 mM tris HCl pH 6.9, 10 mM EDTA and 30 mM sodium citrate [33]). Uninjected controls were also used.

To identify *Galleria* phagocytes, pHrodo™ Red Zymosan Bioparticles™ (ThermoFisher Scientific) were resuspended in IPS to a concentration of 1 mg/mL. *Galleria* larvae were injected with 10 μL of this solution as outlined above.

### *E. coli* culturing

Both the *E. coli* MG1655 control strain and mCherry strain were provided as a kind gift from Dr Remy Chait (University of Exeter). Both strains were grown overnight at 37 °C with vigorous shaking (230 rpm) in LB broth. Following culture, the bacterial cell suspension was centrifuged at 12,000 rpm for 1 minute in pre-weighed Eppendorf tubes. The resultant supernatant was discarded, and the tube re-weighed to calculate the mass of the bacterial cell pellet. The pellet was then resuspended in the appropriate volume of IPS to a final concentration of 1, 5 or 10 mg/mL of *E. coli* for inoculation. For *E. coli* infections, *Galleria* were injected with 10 μL of the 10 mg/mL bacterial suspension.

### Candida albicans culturing

Initial live *C. albicans* SC5314 [34] cultures were provided by Dr Arnab Pradhan. To induce β-glucan masking, overnight cultures were grown at 30 °C at 200 rpm in Yeast Nitrogen Base without amino acids (Merck) prepared according to the manufacturer’s instructions, containing either 2% glucose, 2% glucose + 2% D/L-lactate (Merck), or 2% D/L-lactate alone [31].

### pHrodo™ conjugation

Both *E. coli* and *C. albicans* were fixed for 20 minutes in 4% Paraformaldehyde diluted in PBS at room temperature with gentle agitation. After fixing, the bacteria/fungi were pelleted by centrifugation at 12,000 rpm for 1 minute and the fixative removed. The pellet was washed and resuspended in PBS to remove residual fixative. This washing was repeated thrice. pHrodo™ Red succinimidyl ester (ThermoFisher Scientific) was resuspended in DMSO and conjugated to the fixed pathogens according to the manufacturers instructions. Once the conjugation was completed, the samples were resuspended in IPS for injection into *Galleria* larvae. Concentrations were based on pellet weight as previously described for live *E. coli*.

### Hemolymph collection

100 μL of IPS supplemented with 1 mM phenolthiourea (PTU - Merck) in Eppendorfs was pre-chilled on ice prior to the start of hemolymph collection. *Galleria* larvae were removed from experimental petridishes and were held firmly and close to the lid of a clean petridish for hemolymph collection. The cuticle was pierced laterally on the larval thorax using a pointed scalpel blade, and the hemolymph was allowed to pool onto the petridish lid following gentle squeezing of the insect body. Once spent, the larval carcass was discarded and the hemolymph was immediately collected using a pipette and placed into the pre-chilled IPS + 1 mM PTU solution. The hemolymph was mixed into the IPS solution by gentle pipetting, and the sample was placed back on ice when sufficiently dispersed. Between 3-5 larvae were bled and collected into a single sample tube depending on the experiment.

### Hemolymph preparation for flow cytometry

To prepare hemolymph samples for analysis by flow, the entire collected sample was passed through a 50 μm CellTrics filter (WolfLabs Ltd) into a 5 mL round bottom tube. The filter was washed through with an extra 800 μL of IPS + 1 mM PTU solution, and the tube and filter were gently tapped on the work surface to encourage flow through where needed, and the sample was placed on ice. A 400 μL aliquot of each sample was placed into a second 5 mL tube for staining with CellMask™ Green (CMG, ThermoFisher Scientific) at 1:2000 and/or DAPI (at 1:1000, ready-made solution, Merck).

### Flow cytometry analysis

All analysis was carried out using an Attune NxT Flow Cytometer (Thermo Fisher Scientific). For each starting sample, both stained and unstained aliquots were analysed. A total of 10,000 events within the user-defined cell gate per sample were collected, and the sample was analysed at a flow rate of 100 μl/min. FCS files were exported to FlowJo (BD Biosciences) for subsequent interrogation and plot visualisation. Data values were imported into GraphPad 9 (Prism) for statistical analysis and graphical representation.

### Confocal imaging of *ex vivo* hemocytes

Hemocytes from pHrodo zymosan injected *Galleria* were collected as described above for hemolymph collection, into 500 μL of IPS + 1 mM PTU solution. For observation by confocal microscopy, hemocytes were adhered to glass coverslips by centrifugation of the hemolymph samples using 6 well plates to house and spin the samples and coverslips. Plates were spun at 500 xg for 10 minutes at 4 °C using a centrifuge with plate adapters. An additional IPS + 1 mM PTU solution was gently added after spinning to the side of the well to wash away any unattached cells.

Prior to imaging, coverslips were removed from wells and placed cell side down onto a microscope slide and secured in place using nail polish. Samples were immediately imaged on a Zeiss LSM 880 AiryScanner confocal using a x40 lens (NA 1.3). Z-stacks were acquired through the depth of the adhered cells at a step size of 0.5 μm. Images were exported to FIJI for analysis and formatting.

### Statistical analyses

For statistical analysis, data were imported into GraphPad Prism 9. ANOVA and t-tests were completed dependent upon the experimental set up. For all tests, ns indicates not significant (p>0.05), * indicates p<0.05, ** p<0.001, *** p<0.005 and **** p<0.0001) respectively. GraphPad Prism was also used to generate all graphs used in figures throughout.

### Figures

Experimental diagrams were created using BioRender.com. Diagrams, graphs and flow plots were imported into Adobe Photoshop for figure building.

## Results

### Development of a *Galleria* live hemocyte flow cytometry pipeline

To investigate whether *Galleria* hemolymph centrifugation has any adverse effects on hemocyte cell profiles observed by flow cytometry, a hemolymph collection pipeline was developed (Figure 1A). Briefly, after hemolymph collection, the sample was filtered, before being split into two aliquots. The first – the ‘filtered’ sample – was then immediately analysed, whilst the second was centrifuged at 500 xg for 5 minutes before being resuspended in IPS to form the ‘centrifuged’ sample. By comparing aliquots from a single starting sample, differences in flow plot profiles could be attributed to post-collection processing, rather than sample variation.

**Figure 1:**
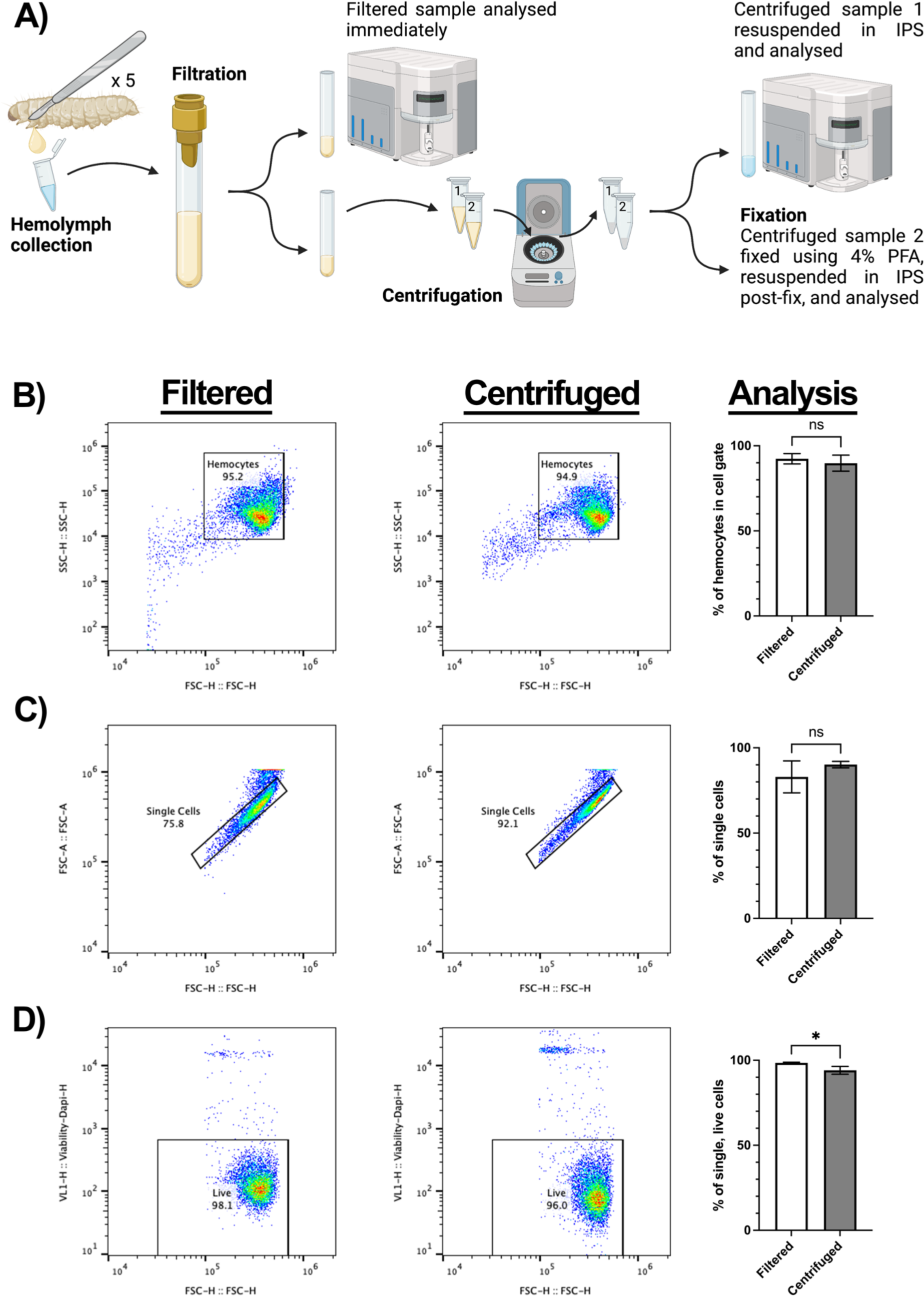
Collection method and analysis of Galleria hemocytes by flow cytometry. A) schematic to show collection and processing methods of hemolymph collected from 5 *Galleria* last instar larvae into a single sample. Samples were split two ways for analysis – filtrated hemolymph only and centrifuged hemolymph. Created using BioRender.com. B) Comparative SSC vs FSC plots for filtered and centrifuged hemolymph. Centrifugation of hemolymph does not significantly concentrate hemocytes into the user-defined cell gate. C) Gating strategy of single cells via visualisation of forward scatter height (FSC-H) to forward scatter area (FSC-A), comparing filtered and centrifuged samples. Centrifugation has no significant effect on the number of single cells in the sample. D) The number of live cells in each sample was determined by the inclusion of DAPI in the hemolymph sample. Centrifugation was found to significantly increase (*, p<0.05, n=3 samples, each containing 5 larvae, unpaired t test) the number of dead cells within the sample.

Initial interpretation of the Forward and Side Scatter (FSC/SSC) plots from both filtered and centrifuged samples revealed a single area of density corresponding to the hemocytes within the sample. Thus, it appears that, in contrast to reports using fixed *Galleria* hemocytes [28,29], live *Galleria* hemocytes are not separable into sub populations based upon relative size and internal complexity alone.

Further analysis comparing post-collection processing methods revealed that the centrifugation and resuspension did not sufficiently remove debris (events outside of the hemocyte cell gate) from the sample or cause changes to the number of single cell events detected (Figure 1B & C). However, the inclusion of DAPI to detect hemocyte cell death revealed that centrifugation causes a small, but significant, reduction in the number of single, living cells in the sample (Figure 1D). Together, Figure 1B-D illustrate the gating strategy employed for all subsequent experiments.

As distinct hemocyte subpopulations have previously been found by flow cytometry when fixed hemocyte samples are analysed [28,29], we repeated our protocol with the inclusion of 4% paraformaldehyde, following either the filtration or centrifugation steps (PFA, Figure 1A). Though there were no discernible differences in the hemocyte cell gate, a less dense secondary population did appear in both the centrifuged and fixed samples (Figure 2A). Using a stricter gating strategy, the secondary population could be gated separately from the main cell population, and the gate was superimposed across all samples to analyse the events within.

**Figure 2:**
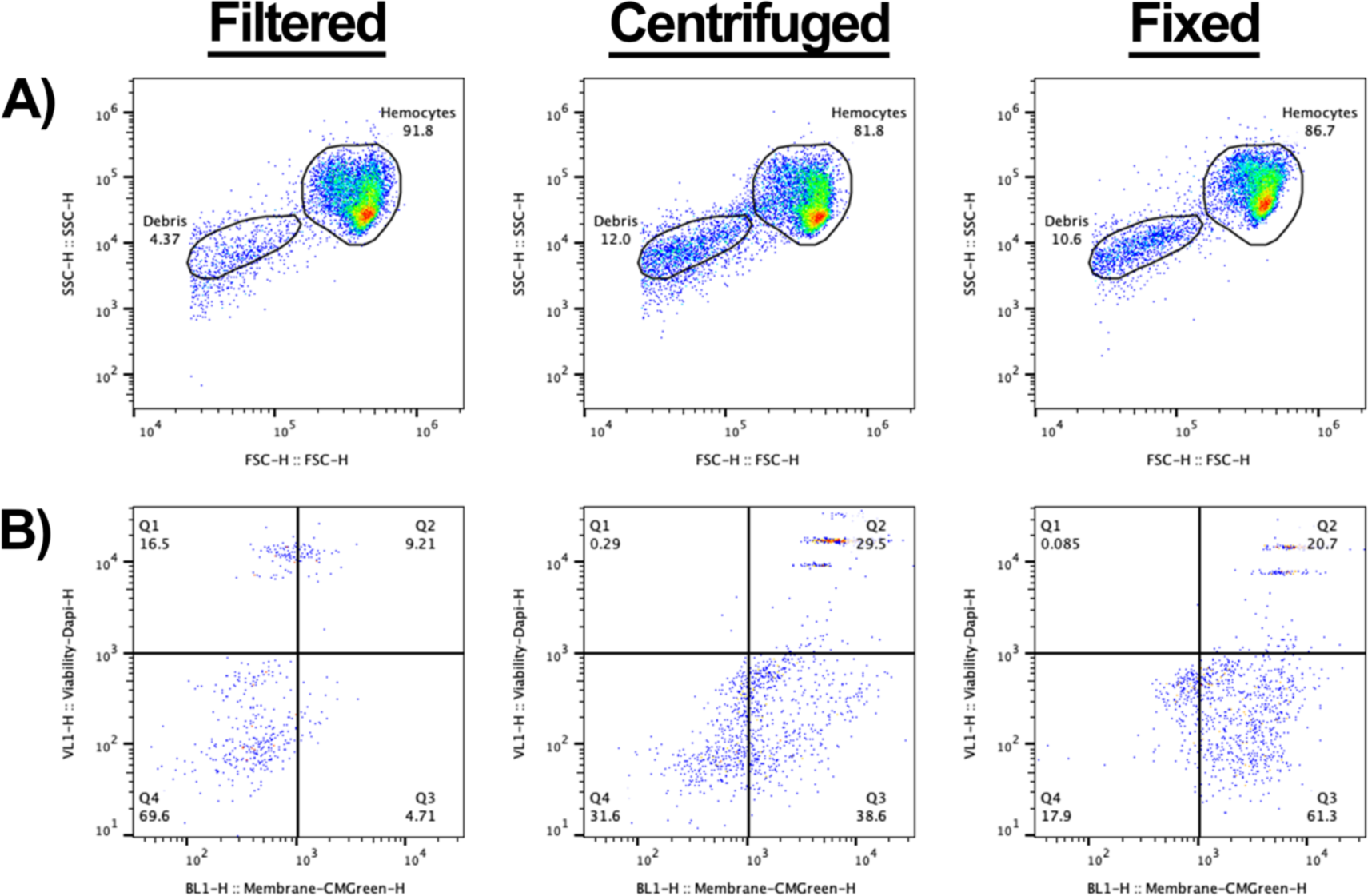
Analysis of non-cellular events within *Galleria* hemolymph. A) Scatter to demonstrate the increase of non-cellular events following centrifugation and fixation of splits of the same starting sample. The non-cellular gate is designated ‘debris’. B) Analysis of events in the debris gate using a cell membrane dye – CMG, and DAPI. Q1 represents CMG negative, DAPI positive, Q2 represents CMG positive, DAPI positive – therefore dead cells –, Q3 represents CMGreen positive, DAPI negative – therefore live cells –, and Q4 represents CMG negative, DAPI negative – true, non-cellular ‘debris’. In the filtered only sample, most events (69.6%) in the debris gate are deemed not cells (Q4). The profile dramatically changes following centrifugation – with more events becoming CMG positive (Q2 and Q3).

To determine the origin of the events within the secondary gate, we added a cell membrane dye (CellMask™ Green (CMG)) and DAPI (a non-permeable fluorescent DNA dye) to the samples prior to analysis. In the filtered only sample, non-cellular debris (CMG negative;DAPI negative) comprised the majority of events (Figure 2B – Q4, 69.6%) with the remaining events split between dead whole cells (CMG positive;DAPI positive) (Figure 2B - Q2) and free DNA-derived material; presumably from lysed cells (CMG negative:DAPI positive) (Figure 2B – Q1). As expected, a negligible fraction was CMG positive but DAPI negative – representing live cells that had evaded the primary gating (Figure 2B – Q3).

In relation to the filtered samples, a higher proportion of events in the secondary gate was observed for both the centrifuged and fixed samples (compare Figure 2A with 2A’ and A’’). Moreover, in the centrifuged only sample, the highest proportion of events were found within Q3 (Figure 2B’). As the contents of Q3 are significantly increased following the process of centrifugation when compared to the filtered only sample, we suggest that centrifugation causes the contraction of living cells – which sees them fall out of the primary cell gate (Figure 2A-A’) and into the secondary gate. Finally, the proportion of dead cells within the centrifuged sample also increased in relation to the filtered sample (Figure 2B-B’, Q2), providing further evidence that centrifugation decreases the viability of hemocytes (Figure 1D).

No apparent changes in the hemocyte profile within the cell gate were observed following fixation of the hemocyte sample (Figure 2A’’), in contrast to previous work showing *Galleria* hemocytes cluster based on relative size and internal complexity [28,29]. In the fixed sample, the majority of the events in the secondary fell within the CMG+, Dapi-quartile (Figure 2B’’, Q3). As all fixed cells should readily take up Dapi, these events likely represent burst, anuclear cells. Only 20% of the events within the secondary gate had a cellular signature (Figure 2B’’ – CMG+, Dapi+, Q2) – representing just 2% of the total events observed for the fixed sample.

These results suggest that post-collection processing methods such as centrifugation and fixing decrease the viability of *Galleria* hemocytes isolated from the circulating hemolymph. They also demonstrate that hemocytes form a single cluster based on relative size and internal complexity when analysed by flow cytometry.

### Using live hemocyte flow cytometry to quantify phagocytosis

As the scatter profiles of live *Galleria* hemocytes failed to distinguish sub-populations of innate immune cells, we sought to identify phagocytes based on the uptake of fluorescent particles. To achieve this, we injected pH sensitive zymosan particles (pHrodo™ Red) into last instar larvae at a concentration of 1 mg/ml, recovering the hemolymph after 2, 4 or 24 hrs (Figure 3A). Confocal imaging revealed fluorescent particles within hemocytes after 2 hrs, and 4 hrs, but with very few cells containing fluorescence after 24 hrs (Figure 3B). As the pH sensitive nature of the pHrodo™ dye means it only fluoresces when internalised, due to the lower pH inside the phagolysosomal system, we conclude that a proportion of the hemocytes constitute phagocytes, which are capable of degrading the zymosan particles over 24 hrs.

**Figure 3:**
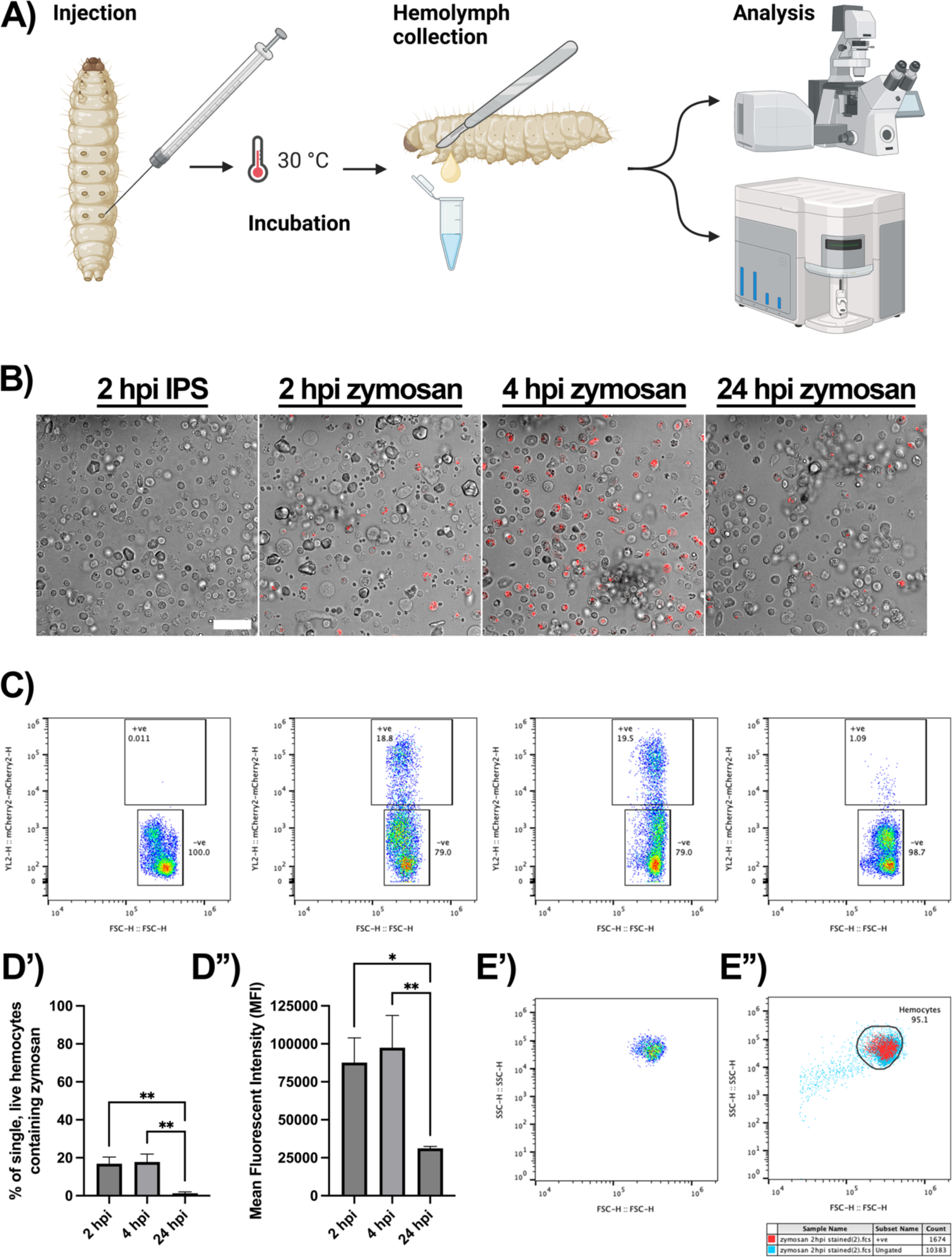
Identifying Galleria phagocytes by the uptake of fluorescent pHrodo™ zymosan particles. A) Schematic to show injection pipeline through to analysis. Following injection with pHrodo*™* zymosan particles, *Galleria* were returned to 30°C (rearing temperature) for incubation, before hemolymph was collected for analysis at set timepoints. Created using BioRender.com. B) Hemocytes from injected larvae visualised by confocal microscopy. Phagocytosed zymosan particles (red) are seen at 2, 4 and 24 hours post injection (hpi). Scale bar at 50 μm. C) Scatter plots to show the proportion of live, single hemocytes containing pHrodo™ zymosan particles based on red fluorescence shift of events. D’) Quantification of phagocytosis over time via flow cytometry. There is a significant reduction in hemocytes containing particles at 24 hpi compared to be 2 and 4 hpi (**, p<0.01, n=3 samples of 3 larvae per sample, one-way ANOVA with multiple comparisons). D’’) Mean fluorescent intensity (MFI, arbitrary units) of events within the zymosan positive gate. MFI significantly decreases by 24 hpi (*, p<0.05 and **, p<0.01, n=3 samples of 3 larvae per sample, one-way ANOVA with multiple comparisons). E’) Representative scatter plot of pHrodo™ positive hemocytes reveals a single cluster of cells based on relative size (FSC) and internal complexity (SSC). E’’) Overlay of pHrodo™ positive hemocytes (red) onto the total hemocyte population (blue) reveals no distinct clustering of phagocytes within the entire hemocyte population.

To further quantify their uptake, we next subjected live hemocyte samples injected with zymosan particles to flow cytometry, according to our filtration protocol (Figure 1A). Following gating for live, single hemocytes, the gated cells were further analysed for the presence of the pHrodo™ signal. The signal could indeed be detected within the hemocytes (Figure 3C), following a similar time-dependent profile to that observed by confocal microscopy. Analysis of the data revealed that the majority of particles were already taken up by 2 hours post injection (hpi), and that they had been predominantly cleared by 24 hpi (Figure 3D’). Moreover, the mean fluorescence intensity (MFI) of the positive cell gate (Figure 3D’’) was significantly decreased at 24 hpi, in relation to 2 hrs, presumably corresponding to the processing and degradation of the particles over time.

As with total hemolymph (Figure 1B) the pHrodo™ positive phagocytes formed a single cluster based on relative size (FSC) and internal complexity (SSC) (Figure 3E’). To determine whether the phagocytic hemocytes clustered within the entire hemocyte population, the pHrodo™ positive cells were overlaid onto the initial FSC-SSC scatter plot. This revealed that the phagocytes did not form an identifiable cluster and were relatively uniform throughout the main hemocyte cluster (Figure 3E’’ – red events represent pHrodo™ cells, blue events represent the total sample). We therefore conclude that functional, live *Galleria* hemocyte populations cannot be characterised based on relative size and internal complexity alone.

### Using live hemocyte flow cytometry to monitor infection dynamics

We next sought to apply our live hemocyte flow pipeline to live bacteria, in order to investigate its efficacy in monitoring infection dynamics. *Escherichia coli (E. coli)* constitutively expressing chromosomally encoded mCherry were injected into last instar larvae, with hemolymph collected, filtered, and subjected to flow cytometry 1,2, 4, 6 or 24 post injection. As with pHrodo™ zymosan, the fluorescent mCherry signal could be identified within single, live hemocytes (Figure 4A).

**Figure 4:**
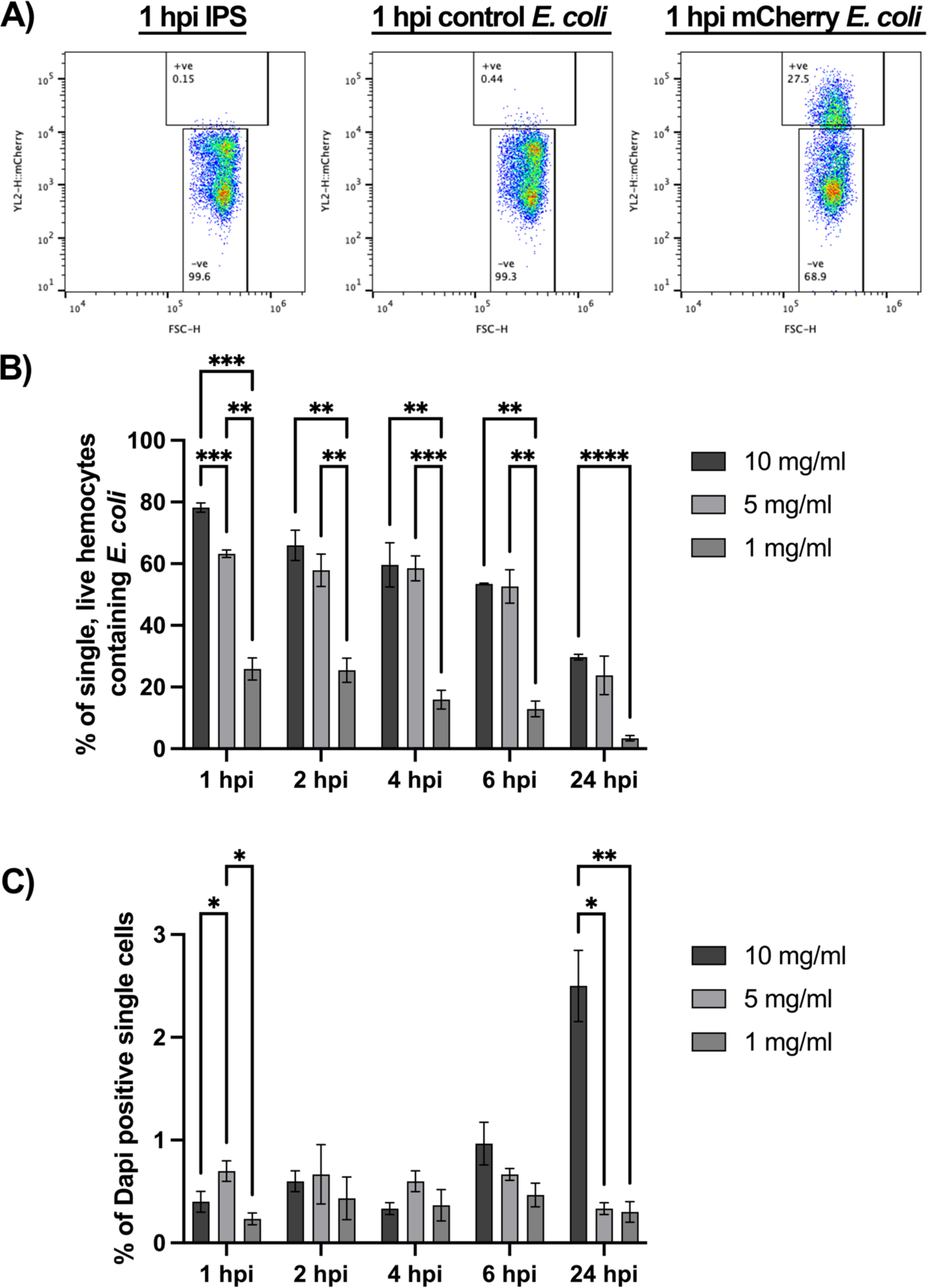
Understanding the response of *Galleria* hemocytes to a live bacterial infection using mCherry *E. coli*. A) Representative scatter plots of single, live hemocytes from IPS control, control *E. coli* and mCherry *E. coli* injected *Galleria* analysed at 1 hour post injection (hpi). Cells from larvae injected with mCherry *E. coli* show an increase in mCherry fluorescence detected (y axis). The shift in mCherry signal is specific to the chromosomally encoded mCherry *E. coli* strain. B) Comparison of mCherry positive hemocytes across different injection doses. C) Analysis of live cells across an infection timecourse and different injection doses. For all graphs statistical significance was determined using 2way ANOVA with Tukey’s multiple comparisons. **** p<0.0001, *** p<0.001, ** p<0.01 and * p<0.05, N=3 samples of 3 larvae each for all doses and timepoints.

To investigate the maximum phagocytic capacity of hemocytes responding to a live bacterial infection, we injected different doses of fluorescent *E. coli* into the larvae, prior to flow cytometry. All injection doses – 1, 5 and 10 mg/mL – showed maximal uptake of fluorescent bacteria at 1 hpi, indicating rapid engulfment of the invading pathogen by the *Galleria* immune cells (Figure 4A and B). Interestingly, while the number of hemocytes containing mCherry *E. coli* increased in a dose-dependent manner at 1 hpi (Figure 4B), this trend was not apparent from 2 hpi onwards in the higher concentration (5 and 10 mg/mL) samples, suggesting processing and degradation of the pathogen occurs within this timeframe. Moreover, although the maximum percentage of live, single hemocytes containing fluorescent *E. coli* increased with increasing dose at 1 hpi, this was not linear; 63.2% (± 1.2 N=3) of hemocytes injected with 5 mg/mL contained fluorescent particles while 78.2% (± 1.5 N=3) of hemocytes injected with 10 mg/mL contained fluorescent bacteria. It is therefore likely that the maximum phagocytic capacity of the *Galleria* immune cell repertoire plateaus at ∼80%.

To investigate whether the *E. coli* strain used influenced hemocyte viability, we quantified the number of DAPI positive single cells in each sample. This revealed dose-dependent significant differences in the number of dead hemocytes at both the 1 and 24 hpi timepoints – with the highest percentage of dead cells (2.5% ± 0.346, N=3) found at 24 hpi in samples from larvae injected with 10 mg/mL *E. coli.* Intra-dose analysis further revealed a significant accumulation of dead cells over time in the 10 mg/mL injection dose at 6 hpi (p<0.001, 2way ANOVA) and 24 hpi (p<0.0001, 2way ANOVA), and a significant reduction in the number of dead cells observed in the 5 mg/mL injection dose when comparing 1 and 24 hpi (p<0.05, 2way ANOVA). At all other time points for all doses, the number of dead hemocytes in the samples remained statistically similar (p>0.05) indicating that hemocytes are rapidly able to deal with non-pathogenic *E. coli* through the phagolysosomal system without significant detriment to the number of live cells within the hemocyte repertoire.

### Using live hemocyte flow cytometry to quantify phagocytosis of fixed fluorescent pathogens

As *Galleria* larvae are used as a host for a wide range of pathogens in microbiological studies, we considered that not all researchers may have an appropriate fluorescent strain of their pathogen of interest. Therefore, we sought to determine whether fixed and labelled pathogens could also be analysed for phagocytic uptake using our developed live cell hemocyte analysis pipeline.

To retain specificity for phagocytic uptake, we purchased the pH sensitive pHrodo dye in a conjugatable form, pHrodo™ Red succinimidyl ester (pHrodo™ Red SE). *E. coli* were fixed with 4% paraformaldehyde, conjugated with pHrodo™ Red, SE, injected into *Galleria* larvae at a concentration of 10 mg/mL and hemolymph extracted at 2 hpi. As with both pHrodo zymosan and live mCherry *E. coli,* a shift in red fluorescence was observed in single, live hemocytes corresponding to the uptake of the fixed conjugated bacteria (Figure 5A). 62.03% (± 2.601, N=3) of single, live hemocytes contained fixed bacteria; similar to the proportion of single, live hemocytes containing live mCherry bacteria at the 2 hpi timepoint following an injection at 10 mg/mL (65.97% ± 4.941, N=3 p<0.2894, unpaired t test) (Figure 4B).

**Figure 5:**
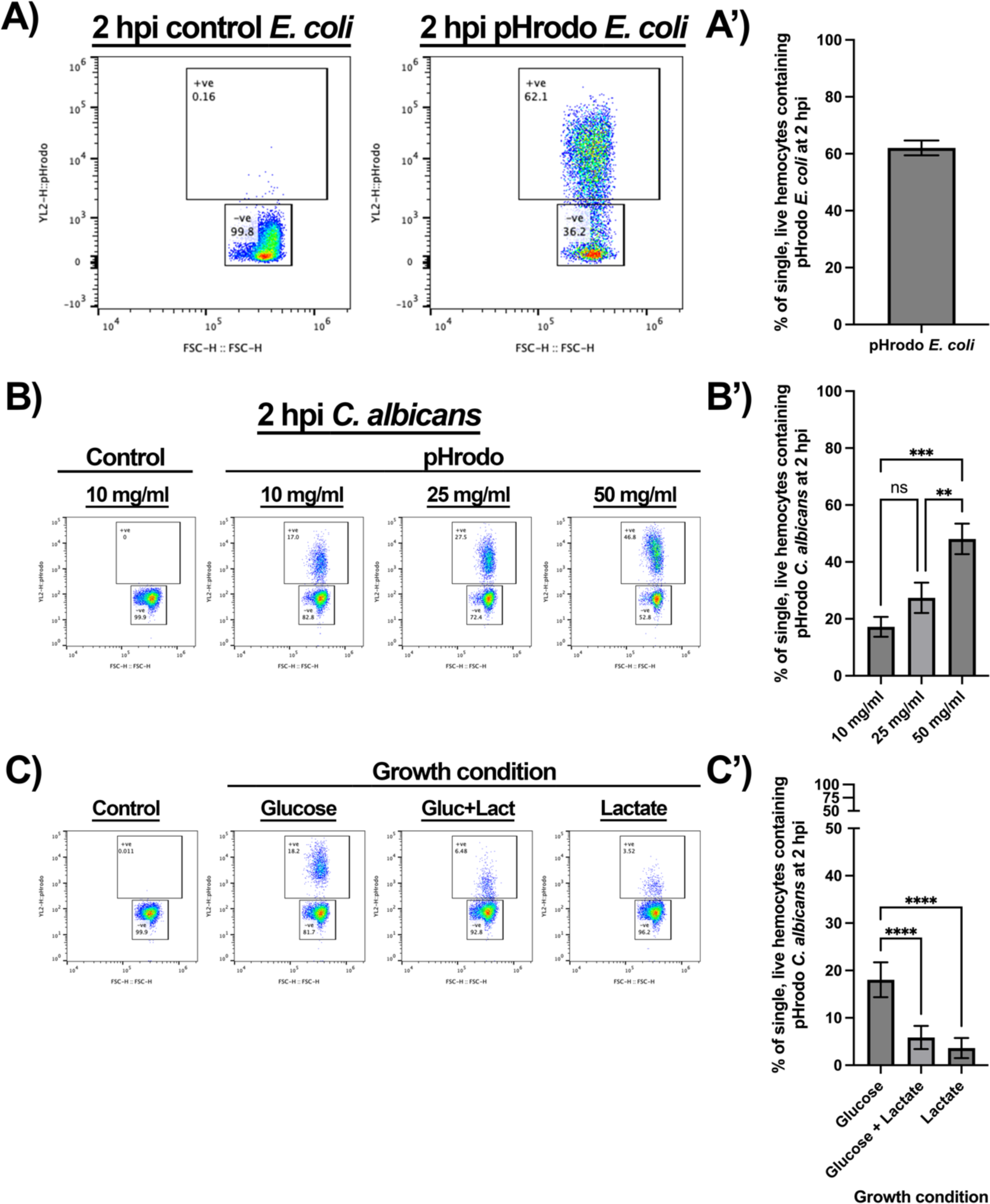
Monitoring fixed pathogen uptake following conjugation of pHrodo™ Red SE. A) Representative flow plots show fluorescence shift following the uptake of pHrodo-*E. coli* by single, live hemocytes at 2 hours post injection (hpi). A’) Quantification of pHrodo-*E. coli* uptake, N=3 samples of 3 larvae each. B) Representative flow plots of hemolymph analysed 2 hpi post fixed pHrodo-*C. albicans* at increasing doses compared to 10 mg/mL control fixed *C. albicans.* B’) Quantification of dose-dependent uptake of fixed pHrodo-*C. albicans*. Significance determined by one-way ANOVA with Tukey’s multiple comparisons, *** p<0.001 and ** p<0.01, N=3 samples containing 3 larvae for each dose. C) Representative flow plots of pHrodo*-C. ablicans* uptake at 2 hpi by single, live hemocytes following growth in the presence of different sugar sources, in comparison to control. C’) Quantification of uptake comparing *C. albicans* growth conditions prior to fixation and conjugation. Significance determined by one-way ANOVA with Tukey’s multiple comparisons. **** p<0.0001, N=11 samples for Glucose and Glucose + Lactate, and N=10 samples for Lactate, each from 3 larvae per sample.

To assess whether this method could be utilised for other types of pathogens, pHrodo™ Red SE was conjugated to the pathogenic fungus *Candida albicans*. As with pHrodo-*E. coli*, uptake of *C. albicans* by *Galleria* hemocytes *in vivo* was indeed observed in a dose dependent manner (Figure 5B and B’). However, even at the highest dose (50 mg/mL), only 48.10% (±5.369, N=3) of hemocytes had engulfed the labelled the fungal cells, whereas ∼80% of hemocytes were able to engulf *E. coli* (Figure 4B). This was possibly due to the larger size of *C. albicans* yeast cells (volume approximately 90 μm^3^ [35] compared to *E. coli* (approximately 1 μm^3^ [36]).

Finally, we sought to determine whether our method could detect known differences in phagocytic uptake. The recognition of the major pathogen-associated molecular pattern, β-glucan, by pattern recognition receptors, such as dectin-1, is critical for phagocytic uptake and antifungal immunity in mammals [38]. However, *C. albicans* modulates immune recognition by masking β-glucan that has become exposed at the fungal cell surface in response to specific environmental signals, including lactate [31,40]. Thus, we exploited the process of β-glucan masking by *C. albicans* to test whether hemocytes, like mammalian macrophages, also display changes in phagocytic uptake in response to β-glucan masking [31,37,39]. Briefly, *C. albicans* SC5314 cells were grown in the presence or absence of glucose and lactate, before being fixed and conjugated to pHrodo™ Red SE. After conjugation, these differentially adapted *C. albicans* cells were injected into *Galleria* larvae at a concentration of 10 mg/mL. At 2 hpi, the hemolymph was extracted and the hemocytes analysed for phagocytic uptake of pHrodo-*C. albicans.* This revealed a significant decrease in phagocytosis of the β-glucan masked *C. albicans* cells grown in the presence of lactate, regardless of the inclusion of glucose (Figure 5C and C’). This differential phagocytic uptake of masked and unmasked *C. albicans* cells faithfully replicates *in vitro* data using murine macrophages [39], thus demonstrating the reliability of the developed assay as well as the relevance of the *Galleria* infection model.

## Discussion

Over the last 20 years the greater waxmoth, *Galleria mellonella* has increased in popularity as the replacement and complementary model organism of choice for *in vivo* infections experiments in place of mammals. *Galleria* larvae have advantages over other non-mammalian models, such as *Drosophila* or zebrafish larvae due to their relatively large size – which makes them easy to handle and dose – and the ability to raise them healthily at human physiological temperature.

The understanding of the immune cell repertoire of *Galleria* larvae dates back over 50 years [22], and little progress has been made to further characterise hemocyte subtypes, with reports focusing on cells that have been physically manipulated via either centrifugation or fixation. In this work, we sought to characterise live *Galleria* hemocytes by flow cytometry with the aim of quantifying distinct hemocyte subtypes following a minimal processing method in order to best maintain cell integrity and morphology.

In contrast with previous reports demonstrating separable sub-populations of fixed *Galleria* hemocytes based on relative size and internal complexity by flow cytometry [28], we found that live hemocytes, subjected to a minimal processing method, behaved as a single population. Which instead corroborates more recent methods to analyse fixed hemocytes by flow cytometry [27]. Even with the use of 4% PFA (Figure 2A) we failed to observe distinct hemocyte groups based on FSC and SSC. Instead, a single cluster of fixed cells was still observed with an accumulation of debris – with our FSC vs SSC plots showing remarkable similarity to other fixed *Galleria* hemocyte analysis [41], where researchers also gate a single population of cells separately from smaller and less complex events which appear on the flow plots. The simplest explanation for this discrepancy is that the previously described subpopulations of hemocytes [28] could appear due to artefacts from sample processing. This is supported by the observation that one of the assigned cell gates from that study appears in the same vicinity as our described debris gate - which we have conclusively demonstrated does not contain viable hemocytes (Figure 2 A and B). Further to this, others also report that the use of fixatives, as well as altering centrifugation speed, had an overall effect on cell morphology – as shown by clear differences in the FSC vs SSC scatter plots [27]. However, we do not rule out the possibility that the hemocyte landscape changes developmentally, as the larvae progress towards pupation, possibly resulting in a more complex distribution of cells the onset of metamorphosis [23].

As we show here, others have also observed a single hemocyte population by flow when live samples are analysed [26]. This work also reports a high level of cell death following centrifugation of the sample prior to analysis, which confirms our conclusion that filtration alone is preferrable to maximise the number of viable cells for analysis within the hemolymph sample.

Having developed a minimal processing method for hemocyte analysis which maintains maximal cell integrity and viability, we sought to extend the assay to be able to quantify *in vivo* phagocytosis – an immune function of hemocytes that can be directly compared to mammalian immune responses. Fixed-cell flow cytometry has previously been used to measure *ex vivo* phagocytosis by *Galleria* hemocytes [29] where, again, designated hemocyte populations are not apparent in our live cell data. Meanwhile other work conducting *ex vivo* [17,42,43] and *in vivo* [44] phagocytosis assays omit initial FSC/SSC plots and gating strategies – thus comparisons to the work presented here cannot be drawn – other than verifying that fluorescence shift can be detected by flow cytometry in hemocytes that have performed phagocytosis of labelled pathogens.

Using our fluorescent phagocytosis assay, we have concluded that the maximum phagocytic capacity of the entire *Galleria* hemocyte repertoire is up to 80% (Figure 4B). This figure corresponds closely to previous work which calculates that the majority of *Galleria* hemocytes can be classed as either granular cells or plasmatocytes based on phenotypic observations [23] – both of which have further been shown to display phagocytic capability *in vitro* [21,25]. Unfortunately, combining these observations and our fluorescent particle uptake assay did not allow for the resolution of less abundant hemocyte subtypes – oenocytoids and spherulocytes – within the hemolymph by flow cytometry via backgating of the phagocytic cells (Figure 3E’’), which may indicate not only their low abundance, but also their heterogeneity. It is therefore likely that future work to characterise the less abundant hemocyte subtypes by flow cytometry may need to include the use of fixation and antibody staining, until such time that distinct hemocyte fluorescent reporter lines are generated by transgenesis – a method which allows for specific hemocyte subpopulation evaluation in *Drosophila melanogaster* [45].

Though phagocytic immune cell responses are undoubtedly better assayed in response to a live infection, it may often be the case that a fluorescent pathogen of interest is either not easily available or achievable, or that the fluorescence expression is linked to a gene promoter with transient or inducible expression. Through the fixation and pHrodo^TM^ tagging of *C. albicans* for use in our *in vivo Galleria* phagocytosis assay we have demonstrated the suitability for this method in fixed pathogen uptake (Figure 5). Our analysis of insect hemocytes faithfully replicates previously published data showing that *C. albicans* uptake by mammalian phagocytes is strongly influenced by the masking of β-glucan at the fungal cell surface [39]. This is significant because β-glucan masking is thought to represent a means of immune evasion for this major fungal pathogen as it encounters certain host signals in specific niches [47–49].

In conclusion, here we present a rapid method to analyse the immune cell repertoire and assay phagocytosis in *in vivo* partial replacement model – *Galleria* mellonella. We demonstrate that this method is superior to previous attempts to analyse *Galleria* fixed or centrifuged *Galleria* immune cells due to improved cell viability and integrity. We corroborate previous findings which demonstrate that the majority of *Galleria* immune cells display phagocytic capability, and detail how the assay can be used to quantify phagocytosis over time – an immune response which, unlike the melanisation of the organism, is relevant to mammalian immune responses. Moreover, the assay developed here produces statistically significant results with only 30 larvae used per condition – representing a reduction of numbers of a model already used as a mammalian replacement, thus further refining the *Galleria* infection model in line with 3Rs principles [46].

## Acknowledgments

We would like to thank James Pearce and Ivan Canada Luna for their help in maintenance of the Galleria laboratory colony, and for stimulating discussions. We also thank Remy Chait (Biosciences, University of Exeter) for his provision of the E. coli strains used in this work. This work was funded by an NC3Rs Project Grant awarded to JGW and AJPB (NC/T001518/1). AJPB was also funded by grants from the Medical Research Council UK (MR/M026663/2) and Wellcome (224323/Z/21/Z), and AP and AJPB were supported by the Medical Research Council Centre for Medical Mycology (MR/N006364/2).

